# Scrub encroachment promotes biodiversity in wetland restoration under eutrophic conditions

**DOI:** 10.1101/2022.02.24.481733

**Authors:** Ane Kirstine Brunbjerg, Camilla Fløjgaard, Tobias Guldberg Frøslev, Dagmar Kappel Andersen, Hans Henrik Bruun, Lars Dalby, Irina Goldberg, Louise Juhl Lehmann, Jesper Erenskjold Moeslund, Rasmus Ejrnæs

## Abstract

Wetlands are important habitats, often threatened by drainage, eutrophication and suppression of ungulate grazing. In many countries, considerable resources are spent combatting scrub encroachment. Here, we hypothesize that encroachment may benefit biodiversity – especially under eutrophic conditions where asymmetric competition among plants compromises conservation targets.

We studied the effects of scrub cover, nutrient levels and soil moisture on richness of vascular plants, bryophytes, soil fungi and microbes in open and overgrown wetlands. We also tested the effect of encroachment, eutrophication and soil moisture on indicators of conservation value (red-listed species, indicator species and uniqueness).

Plant and bryophyte species richness peaked at low soil fertility, whereas soil fertility promoted soil microbes. Soil fungi responded negatively to increasing soil moisture. Lidar-derived variables reflecting degree of scrub cover had predominantly positive effects on species richness measures.

Conservation value indicators had a negative relationship to soil fertility and a positive to encroachment. For plant indicator species, the negative effect of high nutrient levels was offset by encroachment, supporting our hypothesis of competitive release under shade. The positive effect of soil moisture on indicator species was strong in open habitats only.

Nutrient poor mires and meadows host many rare species and require conservation management by grazing and natural hydrology. On former arable lands, where restoration of infertile conditions is unfeasible, we recommend rewilding with opportunities for encroachment towards semi-open willow scrub and swamp forest, with the prospect of high species richness in bryophytes, fungi and soil microbes and competitive release in the herb layer.

## Introduction

Open fens and meadows are characteristic wetland habitats listed on the EU Habitats Directive and targets for conservation (Council Directive 92/43/EEC 1992). They are species-rich and host large numbers of rare and threatened species (Bedford and Godwin 2003, Wassen et al. 2005, Grootjans et al. 2006, van Diggelen et al. 2006). Since the mid-20^th^ century, 80 % of European wetlands have been degraded or lost due to e.g. encroachment following abandonment of traditional extensive grazing, and eutrophication (Middleton et al. 2006, Joyce 2014, Verhoeven 2014). Scrub encroachment is part of the natural succession process; open habitats grow into late successional forest in the absence of disturbances, such as lightning-ignited fire, flooding and grazing (e.g., White 1979, Van Wieren 1995, Bond et al. 2005). However, because of human interference, natural disturbances have diminished overall (e.g., Scholes and Archer 1997, Middleton et al. 2006, Brunbjerg et al. 2014) and the resulting succession has caused widespread scrub encroachment across habitat types and biomes from savannas and steppes to arctic tundra (Naito and Cairns 2011). In Europe, an increase in vegetation density in the period 2001-2015 has been documented and the vegetation change is likely caused by woody regrowth after abandonment of livestock grazing (Buitenwerf et al. 2018). Likewise, in Denmark, 17 % of the area registered as meadow in 1992 has now undergone encroachment (Levin and Nainggolan 2016) however, most of the historical encroachment has happened in the period 1945-1992 (Finderup Nielsen et al. 2021). The pattern is likely to be the same for fens and meadows. When fens and meadows are overgrown with scrub, they lose their legal EU Habitats Directive protection until the scrub eventually grow into late successional swamp forest, which is also protected by the directive (bog woodland 91D0 or Alluvial forests with *Alnus glutinosa* and *Fraxinus excelsior* 91E0). In Denmark, nearly 27 million € are spent annually on support for livestock grazing and mowing in nature areas to combat encroachment and conserve open habitats (Ministry of Food and Environment 2015). Despite the effort, only approximately 20 % of the semi-natural grasslands, including wet meadows and moors, are currently under active livestock grazing (Ejrnæs et al. 2021). This means that most mires are abandoned and subject to free succession. Besides the abandonment of extant fens and mires, many historical fens and meadows have been actively drained, fertilized and ploughed and are today arable fields and leys. As part of the green transition, a large share of this low-lying farmland is projected to be abandoned and rewetted to avoid further carbon loss from the organic soils. While abandonment from agriculture implies a potential for biodiversity, these areas often have large nutrient pools and strongly modified hydrology due to decades of agricultural use. Eutrophication is a threat to species-rich open meadow plant communities due to severe asymmetric competition among plant species increasing with high soil fertility (Grime 1973, Wassen et al. 2005). However, increased shading from encroachment may be hypothesized to relax competition for light among herbs and reduce the competitive exclusion in the field layer as compared to open meadows. Grazing may also partly counterbalance the negative effects of eutrophication (Brunbjerg et al. 2014), but is unlikely to fully compensate (Ejrnæs et al. 2006). Moreover, the full positive effect of disturbances may depend on restoration of natural hydrology (Kołos and Banaszuk 2013, 2018). The combined effects of nutrients, hydrology and disturbance regimes in restored wetlands are difficult to predict, but recent studies indicate that the transformation from arable fields to wetlands often fails to restore the species-rich vegetation consisting of stress-tolerant forbs and bryophytes characteristic for wetlands (Baumane et al. 2021, Kreyling et al. 2021).

Wetland-restoration success is often evaluated on the basis of plants and birds, while important knowledge obtained from other organism groups, e.g. arthropods and fungi, is ignored. In fact, the diversity of heterotrophic organisms, such as arthropods and fungi, is expected to increase with the structural complexity of vegetation and diversity of carbon sources in ecosystems (Elton 1966, Pihlgren and Lennartsson 2008, Brunbjerg et al. 2017). In a recent large-scale study, presence of a shrub layer was the most important variable explaining variation in species richness of fungi and arthropods (Brunbjerg et al. 2020). Heterotrophic organisms gain from the increased biomass following encroachment, as shrubs provide resources and habitats for a large suite of species including herbivores, pollinators, decomposers and epiphytes (Bruun et al. (in press)).

In this paper we investigate the variation in biodiversity along gradients of soil moisture, soil fertility and scrub cover. We further test the hypothesis that occurrence of indicators of high conservation value can be promoted by allowing encroachment to take place – especially in restored wetlands on highly eutrophic former agricultural soils. We suggest two mechanisms for such a positive effect: a) shrubs and trees provide habitat and food resources for large numbers of heterotrophic species, b) a shrub and tree layer may invoke a competitive release in the herb layer reducing the competitive exclusion of typical wetland plants.

## Methods

As part of the present study, we conducted field inventories at 44 wetland sites. The data collection was designed to supplement data collected in the Biowide project, a nation-wide survey of biodiversity in Denmark (Brunbjerg et al. 2019). Biowide included a total of 130 study sites (40 m × 40 m), of which we included all 58 sites evaluated as moist or wet based on plant species composition and soil moisture measurements. The Biowide sites varied in woody species cover from open vegetation over heterogeneous scrub to closed-canopy forest. The Biowide sites also varied in nutrient status from infertile to fertile soils, but were selected to foremost include natural and semi-natural habitats. The additional 44 sites were chosen to increase data coverage of former arable land, semi-natural meadows and agriculturally improved meadows, as well as different levels of scrub encroachment. Sites were located across Denmark, preferably with a minimum distance of 500 m between sites (except two set of sites, where distances were 252 m and 491 m, Fig. 1a). Each site (40 m × 40 m) consisted of four 20 m x 20 m quadrants, each with a 5 m circular plot in the centre (Fig. 1b).

**Figure 1.**
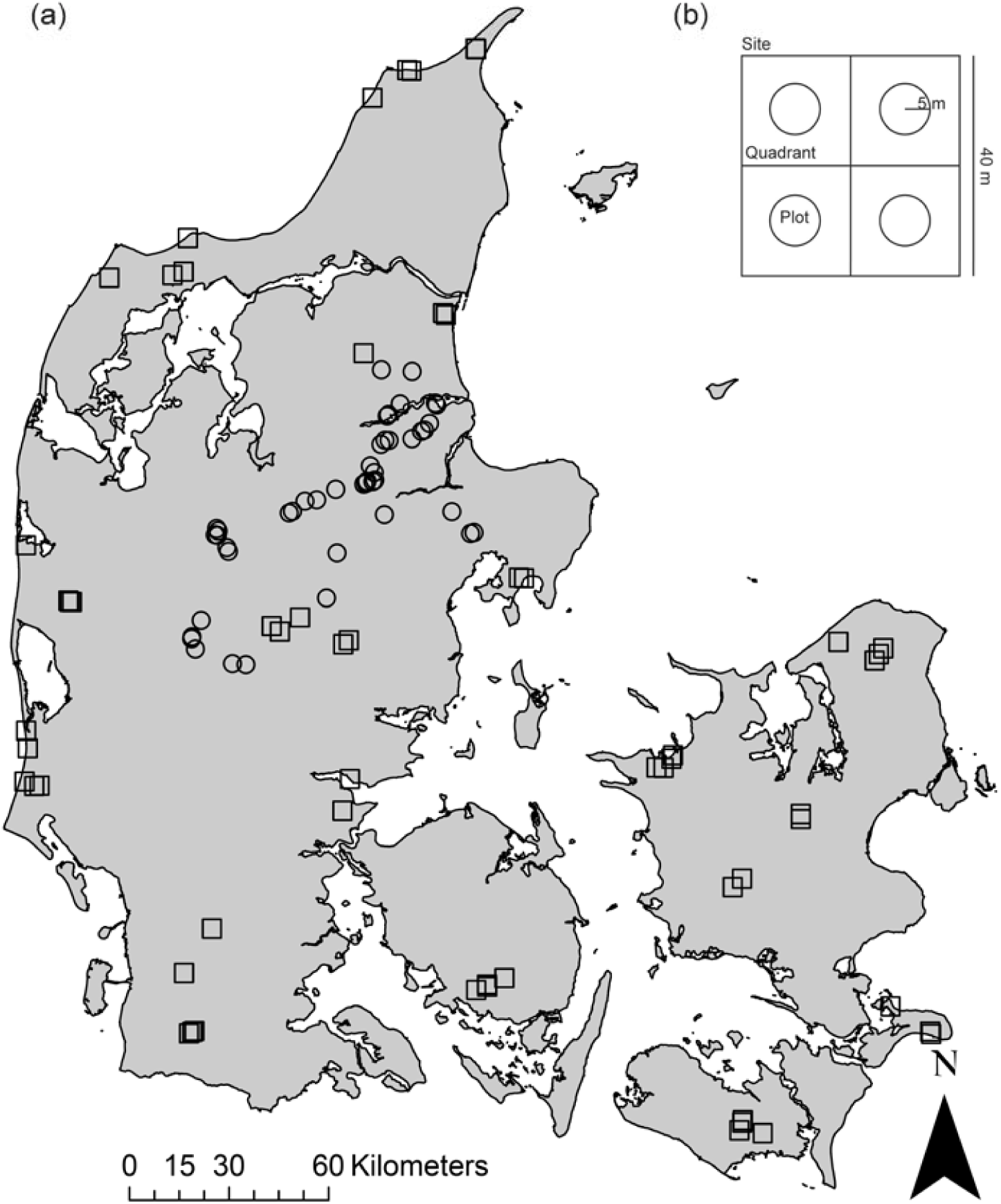
a) Map of Denmark showing the location of the 102 sites (circle: meadow sites, n=44; square: Biowide sites, n=58). b) Site layout with four 20 × 20 m quadrants each containing a 5 m radius circle (plot).

To illustrate the coverage of the soil moisture, nutrient and encroachment gradients covered by the combined data, we compared mean Ellenberg F, N and L values (Ellenberg et al. 1992) for all 5 m circle plots with a reference data set from national monitoring, using identical 5 m circular plots (59,227 sites from agricultural, semi-natural, natural open and forest vegetation, http://www.naturdata.dk) (Nygaard et al. 2017). Mean Ellenberg values were calculated for plots with more than five species present. In scatterplots of plot-mean Ellenberg values, 95 percentile convex hull polygons were drawn for the reference data set as well as the Biowide and meadow data set to visually evaluate the representativity of our data (Appendix A).

### Biowide data collection

Collection of vascular plants and bryophyte data: Vascular plants and bryophytes were inventoried by a trained botanist and exhaustive lists for the four 5 m circle plots were made for each site. In addition, all additional species in the quadrat, but outside the 5 m circles, were recorded. The inventory was done in summer 2014 with a few early spring vascular plant species added in May 2015 (Brunbjerg et al. 2019).

Collection of soil eDNA data: As alternative measures of biodiversity, we used the richness of operational taxonomic units, i.e. OTUs (Blaxter et al. 2005) of fungi and (eukaryotic) soil microbes obtained from metabarcoding of soil-extracted eDNA (Frøslev et al. 2017, Frøslev et al. 2019). We collected soil samples from all sites for the eDNA inventory. Samples were taken in October-November 2014. At each site, we sampled 81 equally distant soil samples from the top c. 15 cm and pooled them after removal of coarse litter. We homogenized the soil by mixing with a drilling machine mounted with a mixing paddle. A subsample of soil for DNA extraction and metabarcoding was taken from the homogenized sample.

### Soil moisture

soil moisture was measured using a FieldScout TDR 300 Soil Moisture Meter. Sixteen equally distanced measurements were taken in each 40 × 40 m site in May 2016 (spring/wet period). To cover the temporal variation in moisture the measurements were repeated in August 2016 (summer/dry period) (Brunbjerg et al. 2020).

### Meadow sites data collection

All additional data collection specifically for the present study was done according to Biowide protocols (Brunbjerg et al. 2019), with the exception of the following: 1) early-spring plants species were not recorded on a separate visit, 2) soil samples were collected during the plant inventory in July-August 2018, i.e. not in November. Soil moisture was measured as in the Biowide project in July-August 2018.

### DNA extraction and metabarcoding

For Biowide and meadow soil samples DNA was extracted and subjected to eDNA metabarcoding through DNA extraction, PCR amplification of genetic marker regions (DNA barcoding regions) and massive parallel sequencing on the Illumina MiSeq platform as described in Brunbjerg et al. (2019). For this study, we used high-throughput sequencing data from marker genes amplified with primers targeting eukaryotes (mainly soil microbes) (18S) and fungi (ITS2). OTU tables were constructed following the overall pipeline in Frøslev et al. (2017). For both fungi and eukaryotes, this consisted of an initial processing with DADA2 (ver. 1.8) (Callahan et al. 2016) to identify exact amplicon sequence variants (ESVs) including removal of chimeras. The preparation of the Biowide-fungal (ITS2) and Biowide-soil microbe (18S) eDNA datasets have been published in Fløjgaard et al. (2019) and Frøslev et al. (2019) respectively, although the fungal dataset was re-sequenced for this study (a detailed description of the sequencing data can be seen in Appendix B).

### Lidar-based measures

In meadows, eutrophication and groundwater level may control the encroachment process and yield different vegetation structures at different combinations of hydrology and eutrophication. Field observations had led us to hypothesize that thickets and woodlands on wet, nutrient poor soils grow more heterogeneous in structure, leaving many canopy openings, compared to thickets and woodlands on more nutrient rich and/or less wet soils. This complexity of vegetation structure can be measured using light detection and ranging (lidar), which is a cost-effective way of gaining fine-resolution data on vegetation structure as compared to field measurements (Lefsky et al. 2002) and which has been shown to capture aspects of vegetation structure that are important and otherwise overlooked for biodiversity (Moeslund et al. 2019b, Thers et al. 2019). A range of variables representing vegetation structure can be derived from lidar, although the translation to and correlation with well-understood properties is not always straightforward. We calculated 23 lidar variables to represent encroachment in the 102 sites. We used the same procedure as in Valdez et al. (2021). The calculations were based on the National near-infrared (1550 nm) lidar-based point cloud (2014-2015) that is freely available from www.dataforsyningen.dk (point density = 4-5 points/m). The final set of variables had a resolution of 1.5 m (except one, see below). For all lidar processing and calculation, we used the OPALS tools (Pfeifer et al. 2014) version 2.3.1 in a Python 2.7 environment. For further details see Valdez et al. (2021). The lidar variables represented both site mean values and standard deviation values to reflect variability within sites. The set of lidar variables encompassed: potential solar radiation (mean and std), adjusted solar radiation (i.e., solar radiation adjusted for vegetation cover; mean and std), amplitude (uncalibrated, but corrected for aircraft type and seasonality, see Valdez et al. 2021), vegetation height (mean and std), vegetation cover (mean, std), mean vegetation density at 0-100 cm, 1 m-3 m, 3 m-10 m and 10 m-50 m, canopy openness (mean, std), terrain openness (mean, std), terrain slope (mean, std), echo ratio (i.e., canopy complexity; mean, std), heat load (std) and mean finescale (0.5m) terrain roughness (Appendix C).

### Response variables

#### OTU richness

As alternative measures of biodiversity, we used the richness of operational taxonomic units, i.e. OTUs (Blaxter et al. 2005) of fungi and soil microbes from metabarcoding of soil eDNA (Frøslev et al. 2017, Frøslev et al. 2019). Classical data collection of fungi is time consuming and OTU richness has been found to resemble classical observed species richness at least for groups of macrofungi that are feasible to include in field inventories (Frøslev et al. 2019). We used OTU richness of fungi and soil microbes as response variables to reflect diversity of species groups not monitored otherwise in this project.

#### Red-listed species

Site richness of red-listed species was calculated for vascular plants and bryophytes based on the current national red list (Moeslund et al. 2019a, Redlist.au.dk).

#### Indicator species

Indicator species include vascular plant species considered moderately to very sensitive to habitat alteration (Fredshavn et al. 2010, see Appendix D). The list of indicator species (Fredshavn et al. 2010) was developed to indicate favorable conservation status according to the Habitats Directive (European Commission 2007). Common to these indicator species is a preference for infertile habitats (low Ellenberg N and high Grime’s S values, Grime 1979).

#### Biotic uniqueness

Uniquity is a scale-dependent metric of biodiversity reflecting how unique the biodiversity of a given site is compared to the gamma diversity across the containing region or collection of sampled sites (Ejrnæs et al. 2018). Uniquity can be calculated based on both observational data as well as non-annotated DNA-data (e.g., OTUs) and hence can reflect both species uniqueness and genetic uniqueness. Contrary to other biodiversity metrics, uniquity accounts for sampling bias and spatial scale. Due to the built-in weighting method, uniqueness of non-annotated DNA-data can be calculated corresponding to e.g. red-listed species (Ejrnæs et al. 2018). Here, we calculated fungi and soil microbe uniquity according to Ejrnæs et al. (2018) in order to reflect the unique site contribution of fungi and soil microbe DNA to the gamma diversity of the collection of sites. Uniquity calculations were based on fungal and soil microbe OUT matrices, site habitat classes and weights from the full Biowide data set (n=130, Brunbjerg et al. 2019) combined with the meadow data set (n=44). The parameter X was set to 1000.

### Explanatory variables

Soil moisture: The trimmed mean of 16 measures per site was used to reflect site soil moisture. For Biowide sites, we used the trimmed mean in August. We detected a systematic discrepancy between moisture in Biowide sites (measured in 2016) and meadow sites (measured in 2018), which could be accounted for by the summer of 2018 being extremely dry. We therefore interpolated the soil moisture trimmed mean values for meadow sites using the predicted values from a k nearest neighbor regression using soil moisture trimmed mean in Biowide sites (n=130, Brunbjerg et al. 2019) as response variable, Ellenberg F values as explanatory variable and k = 11.

#### Soil fertility

Good and reliable field-based measures of nutrient availability are difficult to obtain, as nutrient availability is extremely variable across time and space (Andersen et al. 2013 and references herein). In contrast, the nutrient ratio (mean site Ellenberg N/mean site Ellenberg R, Ellenberg et al. 1992) has been found to reflect eutrophication in wetlands and be highly correlated with the number of typical species in fens (Andersen et al. 2013). For each site we calculated mean Ellenberg N and Ellenberg R values (plant-based bioindication of nutrient status and soil pH, respectively) (Ellenberg et al. 1992). The Ellenberg nutrient ratio was used to reflect eutrophication and the idea of the ratio is to account for the fact that natural nutrient availability in wetlands increases with pH. To avoid circularity in analyses, plant species included in the plantbased conservation indicators (red-listed plants, typical plants) were removed before calculating the nutrient ratios for each model in question.

#### Encroachment

We made a rough classification of sites into two groups (open vegetation and scrub/forest vegetation) based on field photos. The two level factor variable was used as explanatory variable.

### Statistical analyses

#### Richness models

We built generalized linear models with Poisson errors using a set of biodiversity indicators in turn as response variable: vascular plant species richness, bryophyte species richness, fungal OTU richness and soil microbe OTU richness. Soil moisture, nutrient ratio and the 23 lidar-derived variables were used as explanatory variables. Standardized plant species richness was used as covariable in models using OTU richness as response. Negative binomial errors were used if overdispersion was detected in Poisson models (Hilbe 2011). We allowed for interaction between lidar variables and nutrient ratio, and lidar variables and soil moisture, respectively. We included a quadratic term of nutrient ratio and moisture variables if the full model significantly improved according to the ΔAIC < 2 criterion (Burnham and Anderson 2002). The residuals of full models were checked for model misfit and overdispersion and spatial autocorrelation using correlograms from the R package ncf (Bjørnstad 2020). Because of the large number of explanatory variables we used stepwise forward selection using the ΔAIC < 2 criterion (Burnham and Anderson 2002). We used the variation inflation factor (VIF) to test for co-variability among selected explanatory variables and accepted values < 3.

#### Conservation models

In order to test the effect of encroachment on conservation value, we built generalized linear models using a set of biodiversity indicators as response variables: presence of red-listed vascular plants (binomial error), presence of red-listed bryophytes (binomial error), indicator species as defined by Fredshavn et al. (2010) (Poisson error), fungal uniquity and soil microbe uniquity (Gaussian error). Soil moisture and nutrient ratio were used as explanatory variables, and encroachment was tested as a binomial variable discriminating between scrub (woodland with bushes and trees) and open meadows, mires and fens. Negative binomial errors were used if overdispersion was detected in Poisson models (Hilbe 2011). We allowed for interaction between encroachment and nutrient ratio, and encroachment and soil moisture, respectively. We included a quadratic term of the explanatory variables if the full model significantly improved according to the ΔAIC < 2 criterion. The residuals of full models were checked for model misfit and overdispersion and spatial autocorrelation using correlograms from R package ncf (Bjørnstad 2020). We used backwards elimination of explanatory variables using the ΔAIC < 2 criterion (Burnham and Anderson 2002) to reduce full models to final models.

We checked if soil eDNA data for the two datasets could be pooled despite different sampling season (Biowide soil eDNA sampled in November, meadow soil eDNA sampled in July-August), and analyses of similar data showed no marked impact of season on biodiversity measures (fungal and soil microbe OTU richness and uniquity) on the scale where these analyses are concerned.

## Results

The study sites covered the full gradient in nutrient availability of the reference data, but only the wetter part of the reference moisture gradient as expected (Appendix A). The encroachment gradient (Ellenberg L) was also almost fully covered. The meadow data set supplemented Biowide data nicely by adding drier and more nutrient rich sites. Of the 102 sites, we classified 37 as woodland or scrub.

Vascular plant species richness ranged from 12 to 141 species per site. Red-listed plants were found at 57 sites, with a maximum of 18 red-listed species per site. Site bryophyte species richness ranged from 0 to 50 species. Only 21 sites held red-listed bryophytes (1-2 species). Indicator plant species were found in all 102 sites (2-89 species per site).

### Richness models

We found a negative effect of nutrient ratio on plant species richness. However, only c. 7 % of the variation in plant species richness could be accounted for by the model (Table 1). The final model for bryophytes explained 50 % of the variation in bryophyte species richness with a strong negative effect of nutrient ratio and a strong positive effect of vegetation cover on bryophyte richness when looking at effect sizes. The model of soil fungal OTU richness explained c. 45 % of the variation in OTU fungal richness with a strong positive effect of plant species richness. Potential solar radiation and variation in vegetation cover also had a positive effect on fungal OTU richness, while increasing soil moisture, canopy openness and vegetation density at 3-10 m had a negative effects. Potential solar radiation had the largest positive effect on the number of soil microbe OTUs, while fine-scale terrain roughness also affected soil microbe OUT richness positively (Table 1). Nutrient ratio and vegetation density at 3-10 m (impenetrability) interacted, indicating a positive effect of nutrient ratio and a positive effect of encroachment on soil microbe OTU richness except in very eutrophic sites were encroachment had a negative effect (Fig. 2). We found no indication of significant spatial autocorrelation in any of the final models, when checking correlograms.

**Table 1:** Modelling output for GLM negative binomial models using vascular plant species richness, bryophyte species richness, fungal OTU richness and soil microbe OTU richness as response variables and nutrient ratio (Ellenberg N/R), soil moisture and the set of 23 lidar variables as explanatory variables. Estimates, p-values (** < 0.05, *** < 0.01,) and standard errors (in parentheses) are given. DE = deviance explained calculated as (null.deviance-deviance)/null.deviance.

|  | Dependent variable |  |  |  |
| --- | --- | --- | --- | --- |
|  | Plant richness<br>*Negative binomial*<br>DE = 7.3% | Bryophyte richness<br>*Negative binomial*<br>DE = 50.0 % | Fungal OTU<br>richness<br>*Negative binomial*<br>DE = 44.7 % | Soil microbe OTU<br>richness<br>*Negative binomial*<br>DE = 39.9% |
| (intercept) | 3.826*** (0.046) | 2.651*** (0.060) | 5.689*** (0.028) | 5.979*** (0.028) |
| Nutrient ratio | -0.134*** (0.046) | -0.596*** (0.074) |  | 0.096*** (0.031) |
| Soil moisture |  |  | -0.076*** (0.028) |  |
| Plant species richness |  |  | 0.180*** (0.029) |  |
| Vegetation cover |  | 0.672*** (0.080) |  |  |
| Canopy openness |  |  | -0.117*** (0.042) |  |
| Vegetation density 3-10m |  |  | -0.100** (0.041) | 0.076** (0.030) |
| Potential solar radiation |  | 0.169*** (0.060) | 0.118*** (0.029) | 0.115*** (0.027) |
| Complexity (amplitude) |  | -0.171** (0.077) |  |  |
| Fine-scale terrain roughness |  |  |  | 0.074*** (0.028) |
| Vegetation cover variability |  |  | 0.115*** (0.039) |  |
| Nutrient ratio:vegetation density 3-10 m |  |  |  | -0.084** (0.035) |

**Table 2:** Modelling output for models using number of red-listed vascular plants (logistic), red-listed bryophytes (logistic), number of indicator species (GLM negative binomial), fungal uniquity (Gaussian) and soil microbe uniquity (Gaussian) as response variables and nutrient ratio (Ellenberg N/R), soil moisture and encroachment as explanatory variables. Estimates, p-values (** < 0.05, *** < 0.01,) and standard errors (in parentheses) are given. DE = deviance explained calculated as (null.deviance-deviance)/null.deviance. : plant species belonging to the response variable (red-listed plants, typical plants) were removed when calculating the Ellenberg nutrient ratio for each of these model respectively.

|  | Dependent variable |  |  |  |  |
| --- | --- | --- | --- | --- | --- |
|  | Red-listed plants | Red-listed bryophytes | Indicator species | Fungal uniqueness | Soil microbe uniqueness |
|  | *Logistic* | *Logistic* | *Negative binomial* | *Gaussian* | *Gaussian* |
|  | DE=14.7% | DE=17.8% | DE=41.3% | DE=9.5% | DE=16.7% |
| (intercept) | -0.260 (0.221) | -2.835*** (0.581) | 3.078*** (0.066) | 52.577*** (5.218) | 28.424*** (1.849) |
| Nutrient ratio † | -1.020*** (0.254) | -0.804** (0.390) | -0.481*** (0.064) | -14.103*** (4.540) |  |
| Soil moisture |  |  | 0.265*** (0.070) |  | 6.064*** (1.487) |
| Encroachment |  | 2.951*** (0.831) | 0.167 (0.115) | 21.110** (9.396) | 6.552** (3.078) |
| Nutrient ratio:Encroachment |  |  | 0.376*** (0.117) |  |  |
| Soil moisture:Encroachment |  |  | -0.231** (0.100) |  |  |

**Figure 2:**
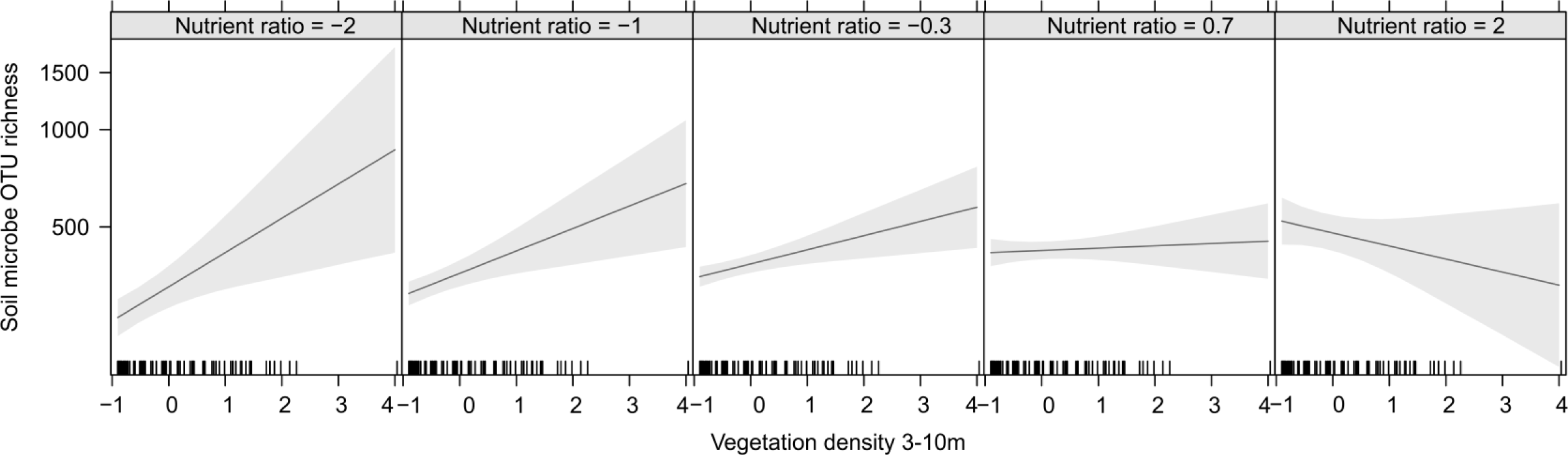
Model output for soil microbe OTU richness illustrating the interaction between nutrient ratio (Ellenberg N/R) and vegetation density at 3-10m. Nutrient ratio and vegetation density are standardized.

### Testing the effect of encroachment on conservation interest

The explanatory strength of conservation models ranged from 9.5 % for the soil fungal uniquity model to 41.3 % for the model for indicator species. Across all response variables, high degree of eutrophication (represented by Ellenberg N/R) affected biodiversity of conservation interest negatively. Encroachment, on the contrary, seemed to have a positive effect on biodiversity indicators. Soil moisture had a positive effect on the number of indicator species and soil microbe uniquity. Interactions between encroachment and nutrient ratio and moisture were only detected in the model for indicator species indicating that encroachment could counterbalance the negative effect of high nutrient levels and dry conditions at least to some degree (Fig. 3). We found no indication of significant spatial autocorrelation in any of the final models when checking correlograms.

**Figure 3:**
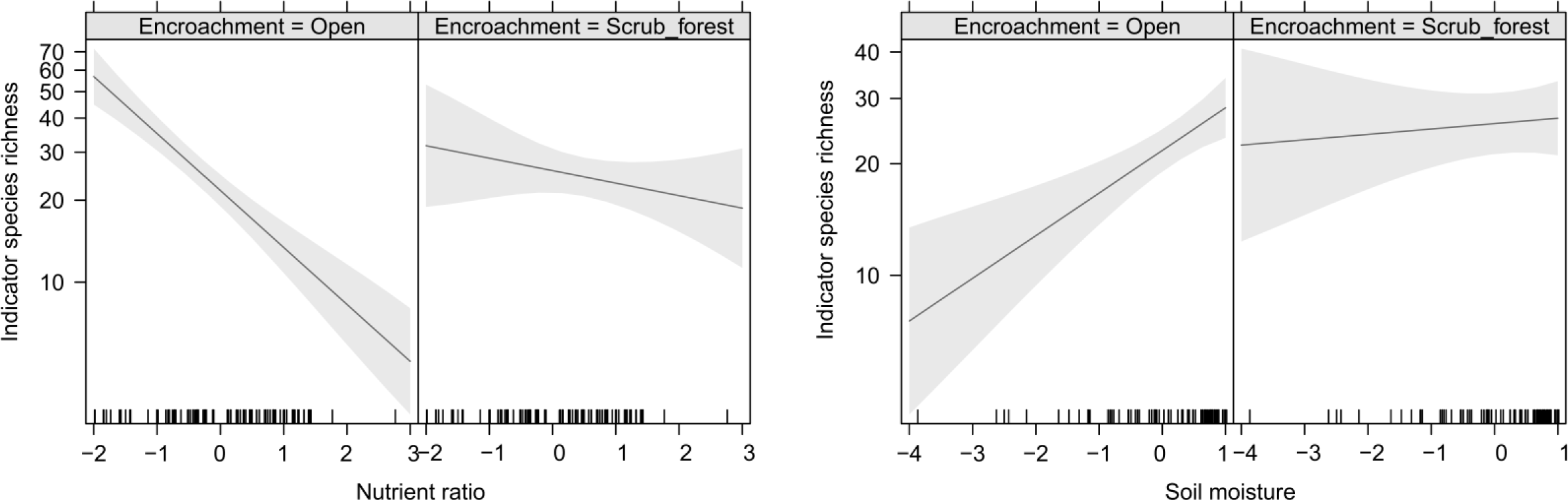
Model output for richness of indicator species illustrating the interaction between a) nutrient ratio (Ellenberg N/R) and encroachment and b) soil moisture and encroachment. Nutrient ratio and soil moisture are standardized.

## Discussion

As expected our study confirmed a rather strong negative effect of soil fertility on the biodiversity of freshwater wetlands, including indicators for conservation status. More surprisingly we found encroachment by shrubs and trees to have a positive effect on red-listed bryophytes, indicator plant species and uniqueness of fungi and soil microbes. Furthermore, we interpret a competitive release after encroachment as the negative response of indicator plants to soil fertility was only present in open wetlands and not in scrub and woodland.

Encroachment is often considered a threat to open-landscape biodiversity (Stoate et al. 2009, Ratajczak et al. 2012). In a conservation management perspective, focus is often on maintaining early successional vegetation, e.g. by grazing and mowing of fens (van Diggelen et al. 2015) to ensure favorable conditions for especially rare plant species sensitive to encroachment (Bart 2021). Traditionally, there has been a focus on vascular plants, when defining habitat types, evaluating conservation status and planning management, e.g. within the framework of the EU Habitats Directive (Brunbjerg et al. 2018). However, this plant-focus may create a biased perception of the effect of encroachment on biodiversity as this effect at least depends on the habitat type and the response group in question (Eldridge et al. 2011). To our surprise, we did not find a negative effect of encroachment in our models for red-listed vascular plants or indicator plants. This is not to say that light-demanding species are not lost through scrub encroachment, but either the loss is relatively weak or losses are off-set by gains of equally rare species. For the indicator plant species the highest richness is observed in infertile and open habitats (Fig. 3) pointing to the need for protecting these against encroachment.

We found a positive effect of encroachment on bryophyte species richness and presence of redlisted bryophytes. Bryophyte richness was higher in sites with relatively high vegetation cover and low lidar amplitude. A high vegetation cover (measured by lidar with 5 points/m^2^) does not necessarily mean that the vegetation is dense (recall that the lidar was recorded during leaves-off and hence it is unlikely to capture the herb layer reliably). Instead, it likely means that bryophyte richness is highest where the vegetation includes small shrubs and trees, which also seem to be the case for the richness of red-listed bryophytes in our study.

Heterotrophic organisms are likely to benefit from the expansion of niches linked to build-up and diversification of organic carbon following encroachment with shrubs and trees (Brändle and Brandl 2001, Bruun et al. (in press)). While the positive effect on bryophytes could be linked to additional substrate for epiphytic species, the beneficial effect of this expansion of ecospace (ss. Brunbjerg et al. 2017) on heterotrophic biodiversity is due to organic plant material for herbivores, symbionts and decomposers as supported by our findings of positive effects of encroachment on fungi and soil microbes. OTU fungal richness showed a complex response with positive effect of canopy closure but negative effect of the density of low trees (i.e., the 3–10 m vegetation layer). In addition there was a positive effect of solar radiation indicating preference for an open park like ecosystem. Only few studies have been conducted successfully linking fungal diversity to lidar variables (but see Thers et al. 2017, Moeslund et al. 2019b) with variables like vegetation structure, successional stage, steep terrain, dead wood and a dense shrub layer being important for fungal species richness. However, the results may not be readily comparable as both of the mentioned studies used macro-fungi from field inventories as response variables instead of OTU fungal richness from soil samples (for a comparison see Frøslev et al. 2019). We could only reproduce the positive effect of tree density on soil microbes under low soil fertility, indicating that in nutrient poor wetlands, trees contribute to ecospace expansion (Brunbjerg et al. 2017), maybe in the form of substrate (i.e., falling leaves or root sap). Lastly, soil microbe richness was higher at relatively high potential radiation and fine-scale terrain roughness, indicating that some abiotic variation in terms of microtopography and vegetation structure promotes richness.

For soil fungi and microbe communities we found more unique assemblages with encroachment, but for indicator species of vascular plants in contrast, we found the classic peak of high species richness in open, nutrient poor fens (Wassen et al. 2005). Others have also found complex richness responses to shrub encroachment, including a hump-shaped relationship (Kesting et al. 2015), indicating that encroachment of scattered shrubs in open grasslands may cause increased habitat heterogeneity which benefit species richness, while complete overgrowth will lead to reduced vascular plant species richness on the scale of small sample plots (Dierschke 2006, Galvánek and Lepš 2008, Ratajczak et al. 2012, Teleki et al. 2020), probably due to light extinction and leaf litter cover inhibiting seedling establishment (Jensen and Schrautzer 1999).

Eutrophication is well-documented as a major threat to freshwater meadow biodiversity. Eutrophication causes a shift in species composition from slow-growing, light demanding vascular plants and bryophytes to more competitive and fast-growing species (Hogg et al. 1995, Bobbink et al. 1998, Bobbink 2004) – a more rapid shift than the vegetation changes due to natural succession (Hogg et al. 1995). Our results are aligned with the negative effect of soil fertility for both species richness of vascular plants and bryophytes, but shows a more complex interaction for soil microbial OTU richness. Hence, the diversity of soil microbes increased with eutrophication and resulting productivity (i.e. available carbon to use as substrate) in open meadow sites. This effect is absent from encroached sites, most likely because shrubs and trees add diverse carbon sources, irrespective of soil fertility. Species of conservation interest and uniqueness of soil fungi similarly showed a negative response to soil fertility. The number of red-listed plant and bryophyte species decreased with increasing soil fertility, as did the unique species assemblages of fungi. Negative effect of eutrophication has been found to be more severe for rare species due to their initial low abundance – at least in grasslands and wetlands (Clark and Tilman 2008).

We hypothesized that in eutrophic sites, competitive release (Keddy and Maclellan 1990) may be a positive consequence of scrub encroachment and the resulting vertical differentiation of vegetation layers. The competitive release hypothesis is underpinned by the notorious depauperate plant species richness in eutrophic herbaceous vegetation due to asymmetric competition for light and nutrients (e.g., Crawley et al. 2005). We found the mentioned strong negative effect of eutrophication on plant species richness in open herb communities, but a much smaller effect under canopy cover for the number of indicator species (Fig. 3), supporting the hypothesis of competitive release.

Rewetting and recreating natural hydrology is a well-established management recommendation for fen and meadow systems (Kołos and Banaszuk 2013, 2018). However, we did not find a general positive effect of soil moisture on species richness and indicators of conservation value in our study, the effect was only positive for indicator species in open habitats but not after encroachment. We suspect that the effect of soil moisture can be partly confounded with both eutrophication and encroachment because leached nutrients in the watershed is transported with the water and released into the wetland communities and also the wettest areas tend to be abandoned first and generally exhibit heavier encroachment than less wet sites.

Anecdotic evidence from our own surveys of aerial photographs indicates that willow scrub and alder swamps predominantly occur in the wettest parts of river valleys, e.g. in places where historical small-scale peat- extraction has left inundated pits and rendered the tract unsuitable for cultivation.

In our study, plant species richness and the number of red-listed plants were solely affected by soil fertility and the models only had limited predictive power (c. 7 % and c. 14 %, respectively). The predictive power of these models may seem low when compared to c. 60 % explained variation in plant species richness in the full Biowide data set in Brunbjerg et al. (2020). Although the two studies are not directly comparable, several explanations may be suggested for this discrepancy e.g. our encroachment variable is rather crude and also variables representing disturbance (e.g., grazing) and historical events related to former cultivation not included in this study could possibly be important for the plant species richness in our wetland sites.

Because of the need for reducing greenhouse gas emissions (the Paris agreement, United Nations 2015) e.g. by agricultural abandonment of organic soils, Denmark is now planning to abandon 100.000 ha of low-lying cultivated areas. Cultivated areas in river valleys are obvious candidates, as these areas often have low agricultural profitability and cultivation results in high carbon emissions (Günther et al. 2020). While setting aside cultivation of these potential wetlands implies a great potential for ecological restoration, our study shows that notably earlier eutrophication caused by decades of arable farming will almost inevitably hamper the restoration target of species-rich meadows and fens. Based on our results, we recommend to combine a relaxed attitude to encroachment with reintroduction of natural disturbances (e.g., widespread rewilding of large herbivores) in order to promote semi-open scrub and woodland communities. Scattered bushes and thickets are natural elements in grazing systems, as many shrub species are vigorous resprouters, e.g. Salix species (Klimkowska et al. 2010). In areas not suitable for year-round grazing, so-called ‘passive rewilding’ (i.e. natural processes are allowed to restore themselves, Svenning et al. 2016) may be a superior solution compared to mechanical harvesting or intensive summer meadow grazing. This strategy for restoration of set aside of former cultivated fields should not supplement and not replace the critical conservation of unique fens and meadows of high conservation value relying on a long an unbroken historical continuity and naturally low nutrient status.

## Author contributions

Ane Kirstine Brunbjerg: Formal analysis, Methodology, Writing - original draft, Writing - review & editing

Camilla Fløjgaard: Conceptualization, Data collection, Formal analysis, Methodology, Writing - original draft, Writing - review & editing

Tobias Guldberg Frøslev: Data collection, Data curation, Writing - review & editing

Dagmar Kappel Andersen: Data collection, Writing - review & editing

Hans Henrik Bruun: Conceptualization, Writing - review & editing

Lars Dalby: Data curation, Writing - review & editing

Irina Goldberg: Data collection, Writing - review & editing

Louise Juhl Lehmann: Data collection, Writing - review & editing

Jesper Erenskjold Moeslund: Data collection, Data curation, Writing - review & editing

Rasmus Ejrnæs: Conceptualization, Formal analysis, Methodology, Writing - original draft, Writing - review & editing

## Appendix A

95 percentile convex hull plots of Ellenberg F, L and N values from a reference data set (http://www.natur_data.dk) of open and forest habitat types (green, n = 59,227) as well as the data sets used in this study: Biowide sites (orange, n = 58), meadow sites (blue, n=44). Black and green dots represent mean Ellenberg values of the 102 sites.

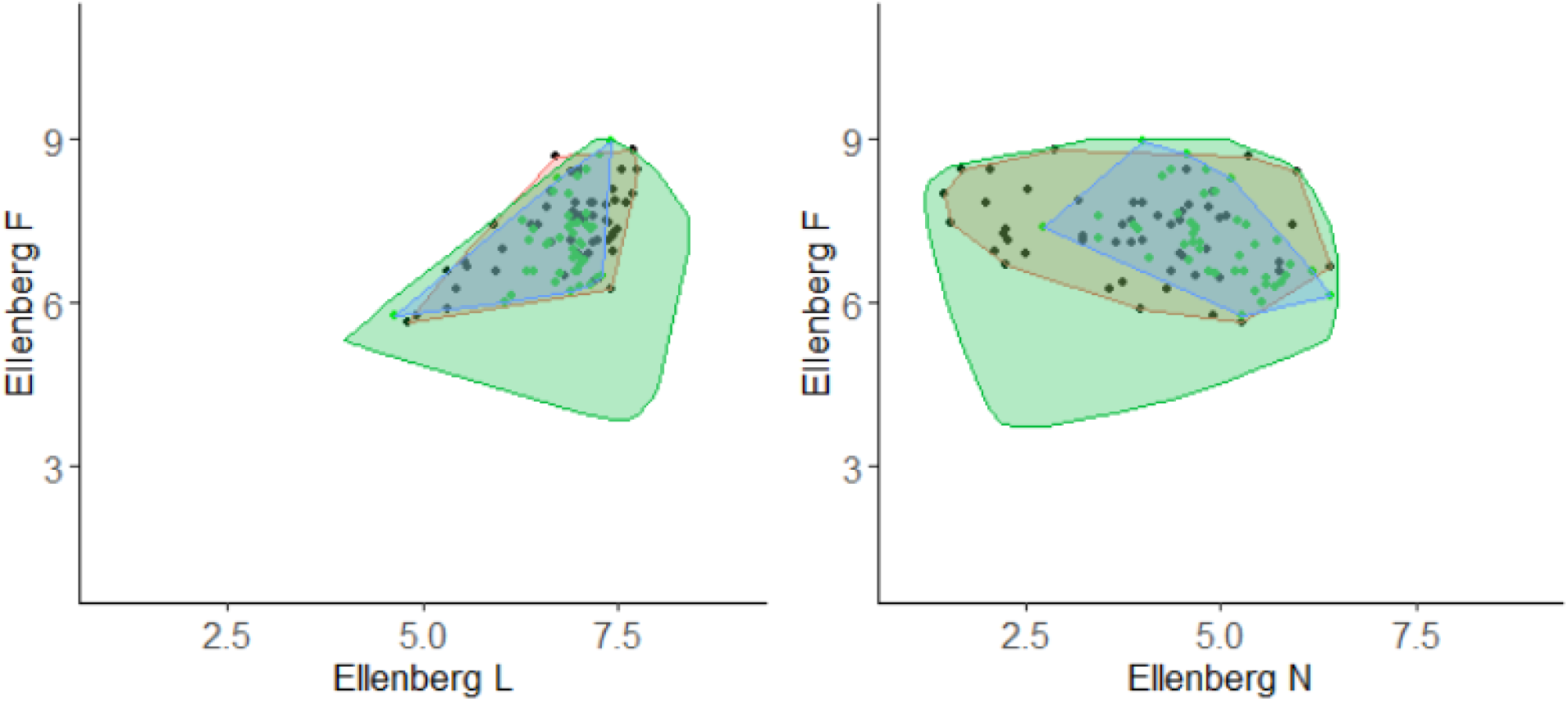

## Appendix B

Detailed information on molecular data

### DNA extraction (44 study sites)

DNA was extracted from 4 g of soil using the PowerMax Soil DNA Isolation kit (Qiagen), following the manufacturer procedure. DNA extract was purified using the PowerClean DNA Clean-Up Kit (Qiagen), and DNA was normalized to 1 ng/μl after fluorometric quantification using the Qubit™ dsDNA HS Assay Kit (Thermo Fischer). The 44 samples were extracted in smaller batches with one negative control for each batch.

### Sequence data

For both the 44 study sites and the 130 Biowide sites, we amplified and sequenced molecular marker regions for fungi and eukaryotes. For fungi the internal transcribed spacer region 2 (nrITS2) was amplified using primers gITS7 (5’-GTGARTCATCGARTCTTTG-3’) (Ihrmark et al. 2012) and ITS4 (5’-TCCTCCGCTTATTGATATGC-3’) (White et al. 1990). For eukaryotes, the primers TAReuk454FWD1 (5’-CCAGCASCYGCGGTAATTCC-3’) and TAReukREV3 (5’-ACTTTCGTTCTTGATYRA-3’) (Stoeck et al. 2010) were used to amplify the V4 rRNA loop of the small nuclear ribosomal subunit (18S nrDNA).

PCR amplifications contained 0.04 U μL-1 AmpliTaq Gold (Life Technologies), 0.6 μM of each primer, 0.8 mg mL-1 bovine serum albumin (BSA) and, 1X Gold Buffer, 2.5 mM of MgCl2, 0.2 mM each of dNTPs and 1 μL (1 ng) DNA extract, in 25 μL reaction volume. For every batch of PCR reactions, three PCR blanks and extraction blanks were included. For fungi was used an initial denaturation step of 5 min at 95 °C, followed by 31 cycles of denaturation of 30 s at 95 °C, 30 s annealing at 55 °C, 60 s elongation at 72 °C, and a final elongation at 72 °C for 7 min. For eukaryotes was used an initial denaturation step of 7 min at 95 °C, followed by first 15 cycles of 30 s at 95 °C, 30 s at 53 °C, 45 s at 72 °C, then 20 cycles of 30 s at 95 °C, 30 s at 48 °C, 45 s at 72 °C, and a final elongation at 72 °C for 10 min. Presence and size of DNA fragments was verified on 2% agarose gel, stained with GelRedTM (Biotium, CA, USA).

Primers were designed with 96 unique tags (MID/barcodes) of 6 bp at the 5’ end, preceded by 1,2 or 3 N’s. No primer tag was used more than once in any sequencing library and no combination of forward and reverse primer tags was reused in the study. PCR products were pooled into pools with approximately the same number of samples, with no tag reused in any pool. Each pool was cleaned with the MinElute purification kit (Qiagen) and built into separate sequencing libraries using the TruSeq DNA PCR-Free Library Preparation Kit (Illumina). This was done following the manufacture procedure, except that all suggested bead purifications were replaced with MinElute purification. Before and after library building, pools were checked on an Agilent BioAnalyzer 2100 to verify the length of the products. Adapter dimers were removed using Agencourt AMPure XP beads, with a 1,5 bead:sample ratio. The libraries were mixed in equal proportions and sequenced together on one 300 paired-end run on the Illumina MiSeq (3000) at the Danish National Sequencing Centre. Eukaryote sequence data for 44 study sites was produced for this study, and bioinformatically combined with data for the 130 Biowide samples which was produced in an earlier study (Fløjgaard et al. 2019). Fungi sequence data for 44 study sites was produced for this study including all above steps, whereas the data for the 130 Biowide sites was produced by resequencing the libraries (from Frøslev et al. 2019) with 300 bp PE sequencing, and bioinformatically combining the two datasets.

### Sequence processing

The bioinformatic processing of the sequence data followed the strategy outlined in Brunbjerg et al. (2019). Demultiplexing of samples was done with custom scripts that keeps R1 and R2 separate for DADA2 processing (Frøslev et al. 2019), and is based on Cutadapt (Martin 2011) – and also Sickle (Joshi and Fass 2011) for fungi – as in Frøslev et al. (2019) searching for a sequence pattern matching the full-length combined tag and primer allowing for no errors, and removing possible remnants of the other primer at the 3’ end. We used DADA2 v 1.8 (Callahan et al. 2016) to identify OTUs (also known as exact amplicon sequence variants, ESVs) and for removal of chimeras (bismeras). For eukaryotes, the OTU tables were used directly. The fungal data was clustered at 98.5 % and filtered to contain only ingroup data – ie. kingdom Fungi. For the fungal dataset, taxonomy was assigned by matching against the UNITE database (Nilsson et al. 2019), and for the eukaryotic data using a custom script based on BLASTn searches against genbank (Altschul et al. 1990).

Sequence data and analytical documentation can be obtained by contacting the first author.

## Appendix C

Lidar-based explanatory variables and interpretation. If the standard deviation of a variable was calculated, in addition to its mean, the variable is denoted with an asterisk.

|  | Explanatory variable | Explanation |
| --- | --- | --- |
| Light/heat | Potential solar radiation* |  |
|  | Adjusted potential solar radiation* | Solar radiation adjusted for vegetation cover |
|  | Heat load index (only standard deviation) | Variation in south-westness |
|  | Canopy openness* | How open the vegetation is (also low vegetation) |
|  | Vegetation cover* | Fraction of vegetation points to ground point |
|  | Terrain slope* |  |
| Microtopography | Terrain openness* | Overall flatness of terrain |
|  | Fine-scale terrain roughness | Fine-scale (0.5m) terrain heterogeneity |
| Encroachment | Vegetation height* |  |
|  | Vegetation density at 0-1 m, 1 m-3 m, 3 m-10 m and 10 m-50 m | Fraction of vegetation points in given layer compared to that in all other vegetation layers |
|  | Amplitude | Succession state (high amplitude = high reflectance/flat and simple vegetation, low amplitude = low reflectance/complex vegetation). The measure is corrected for aircraft type and seasonality, see Valdez et al. 2021) |
|  | Echo ratio* | Canopy cover and complexity |

## Appendix D

Indicator species considered moderately to very sensitive towards habitat changes as defined by Fredshavn et al. 2010. The list of indicator species was developed to indicate favorable conservation status cf. the Habitats Directive (European Commission 2007). Species name and family are given.

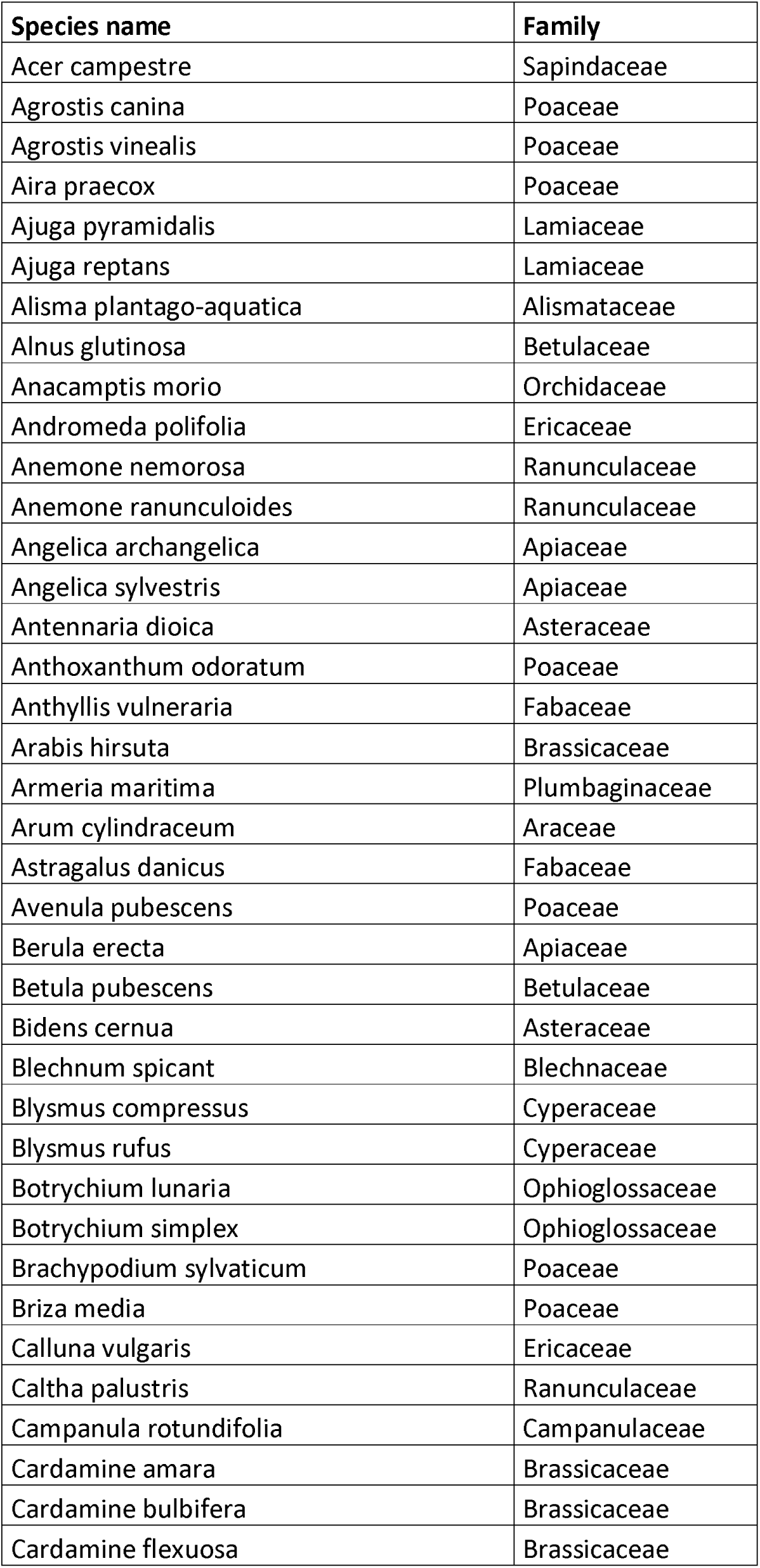
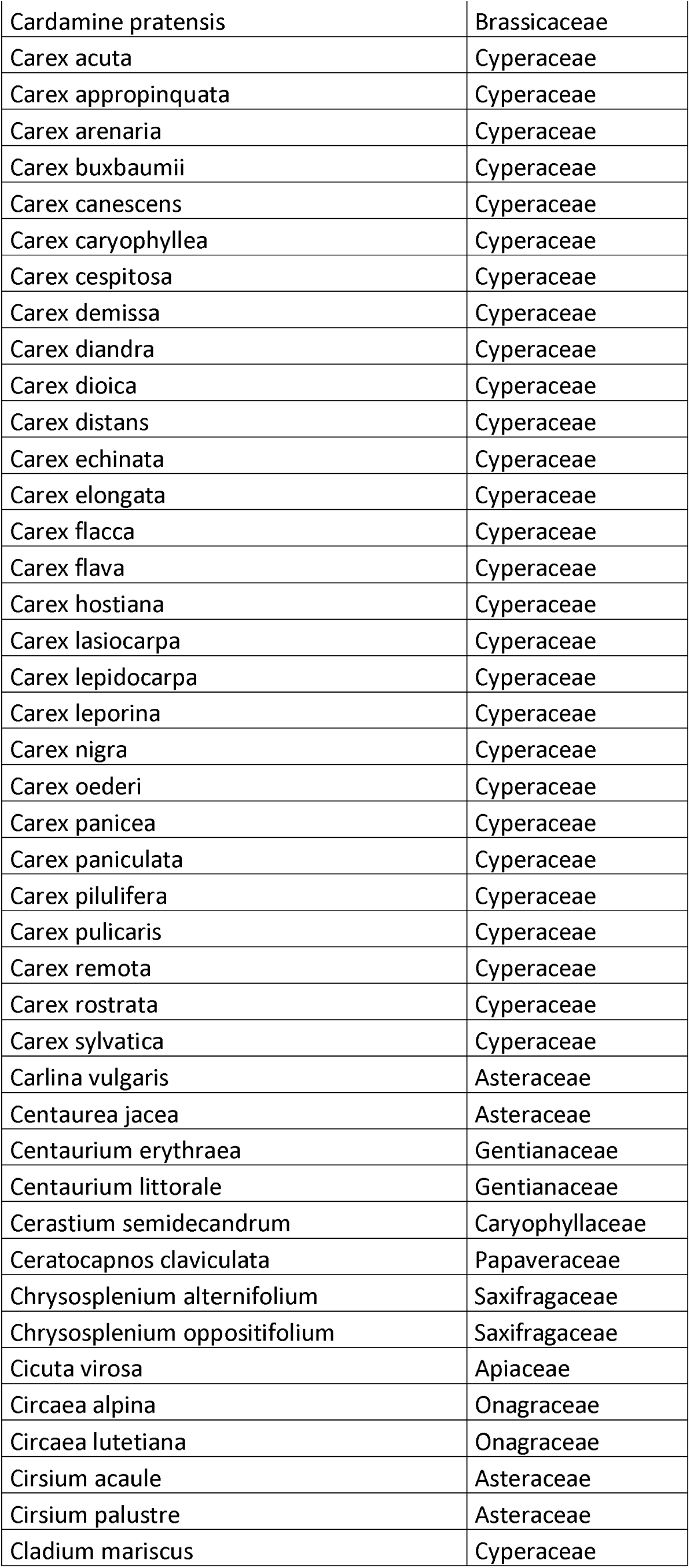
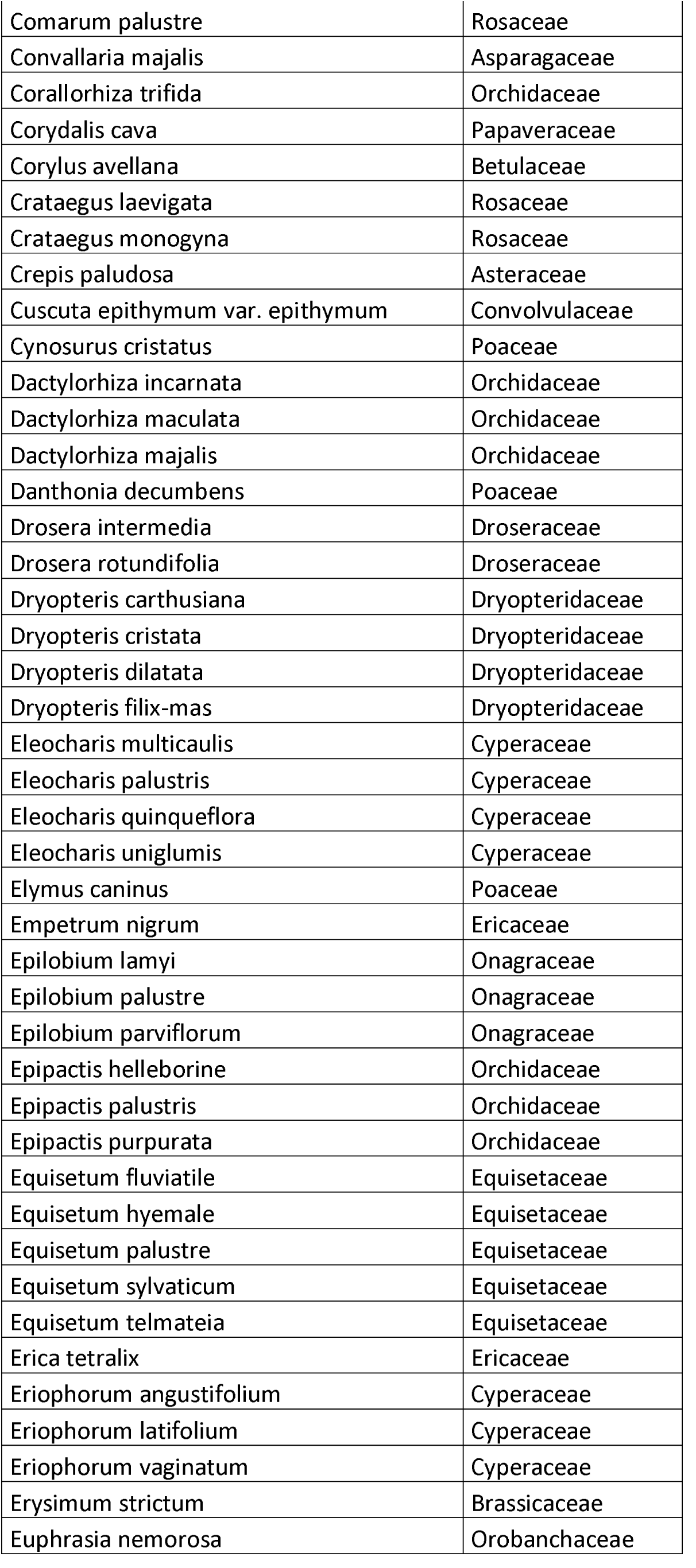
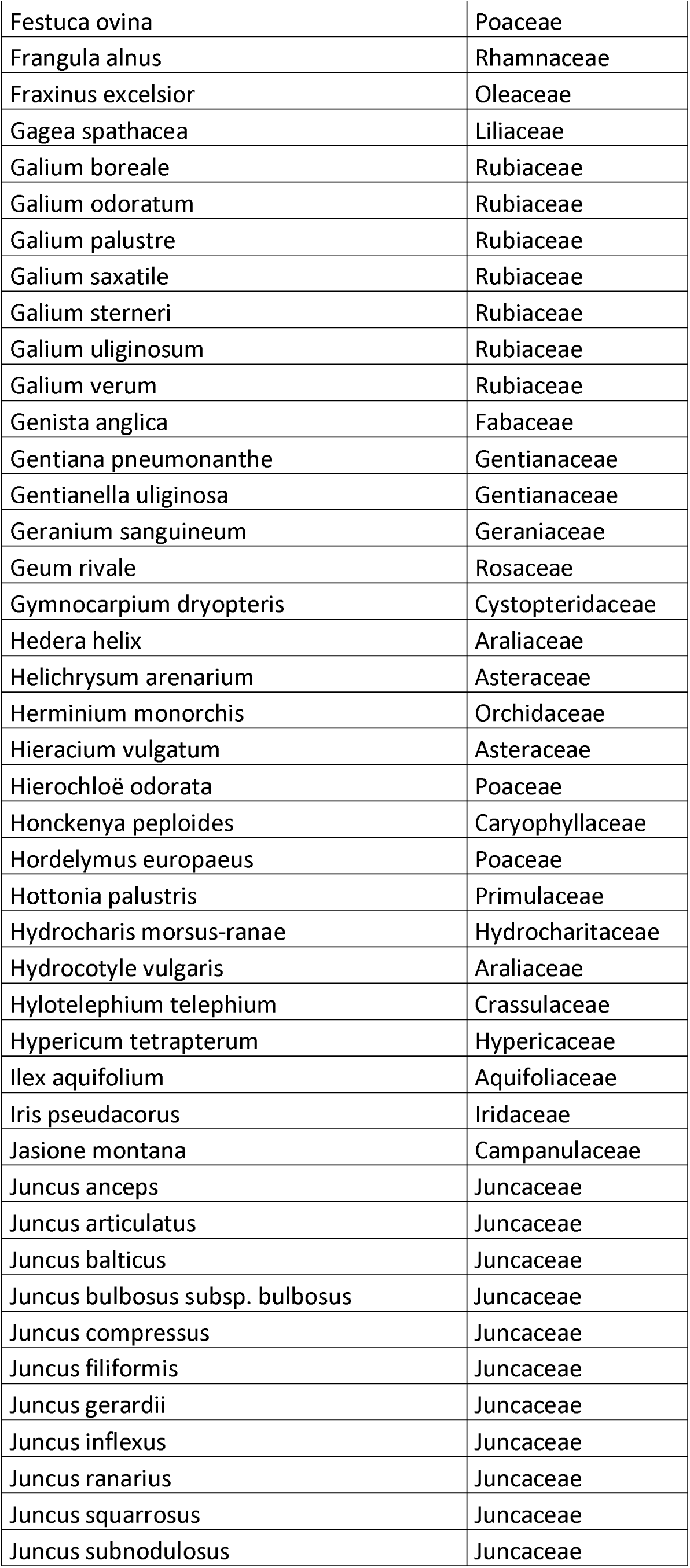
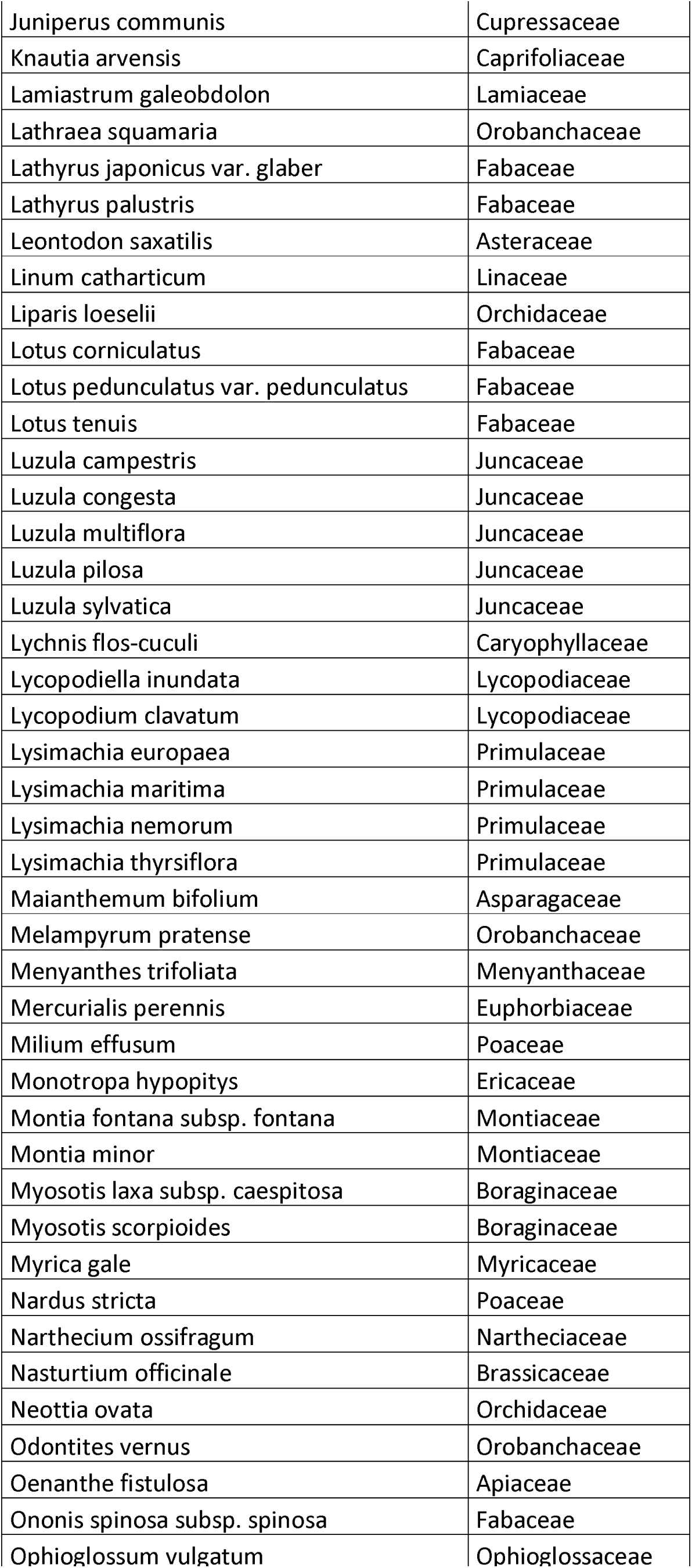
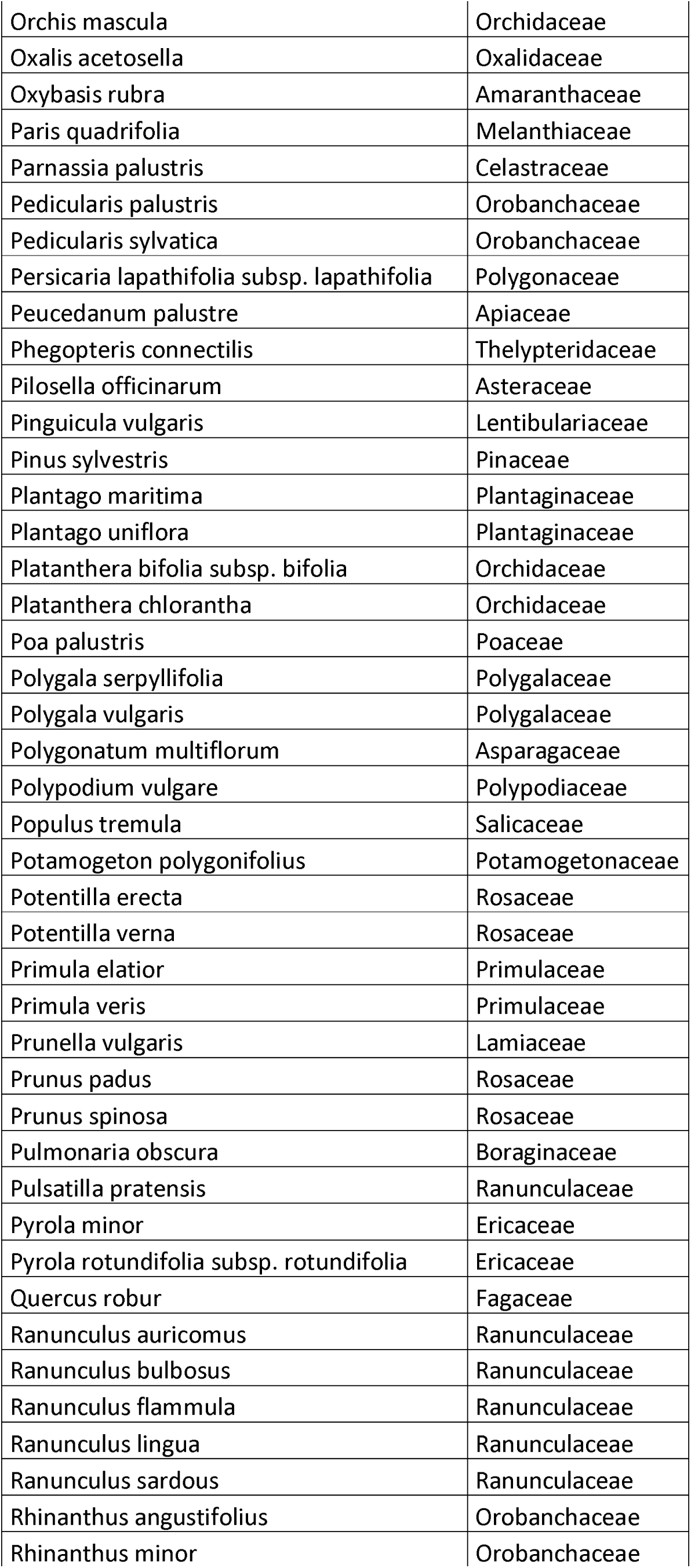
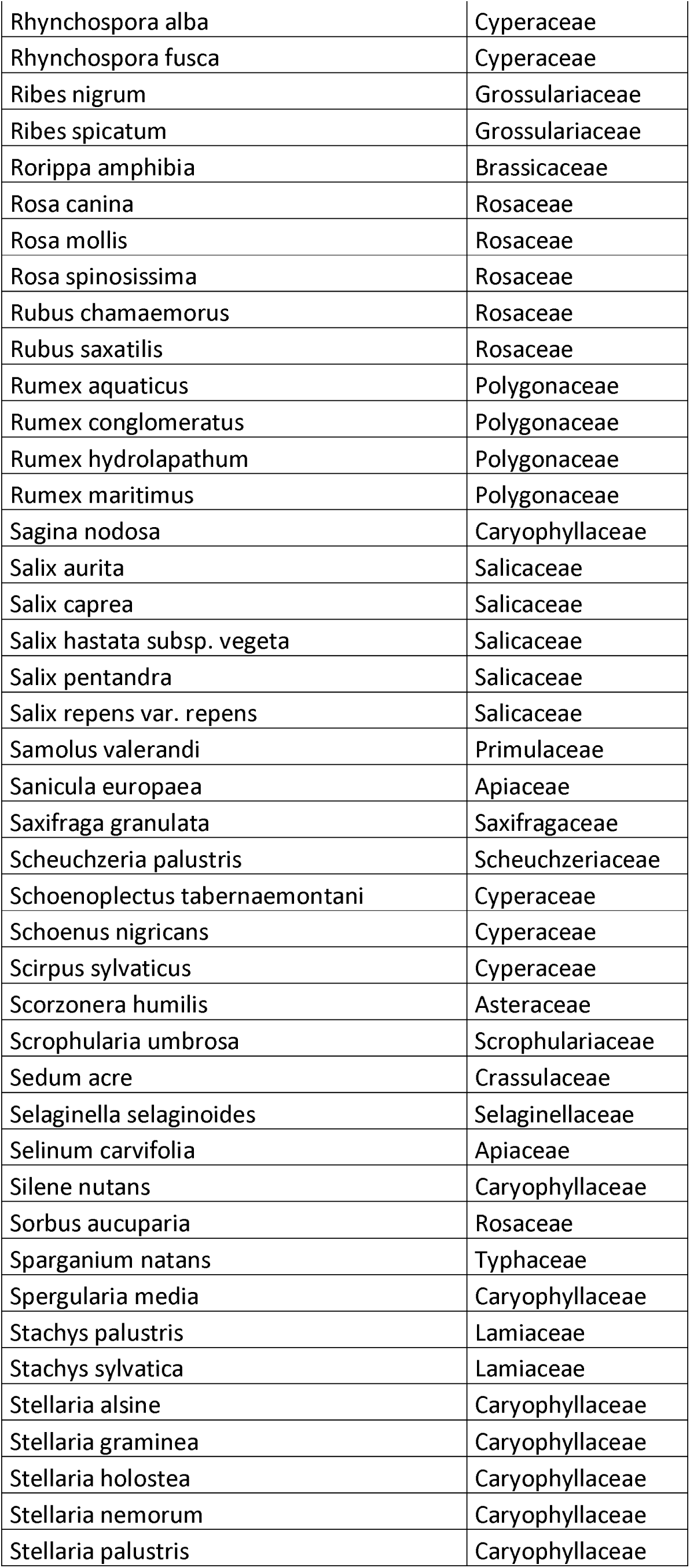
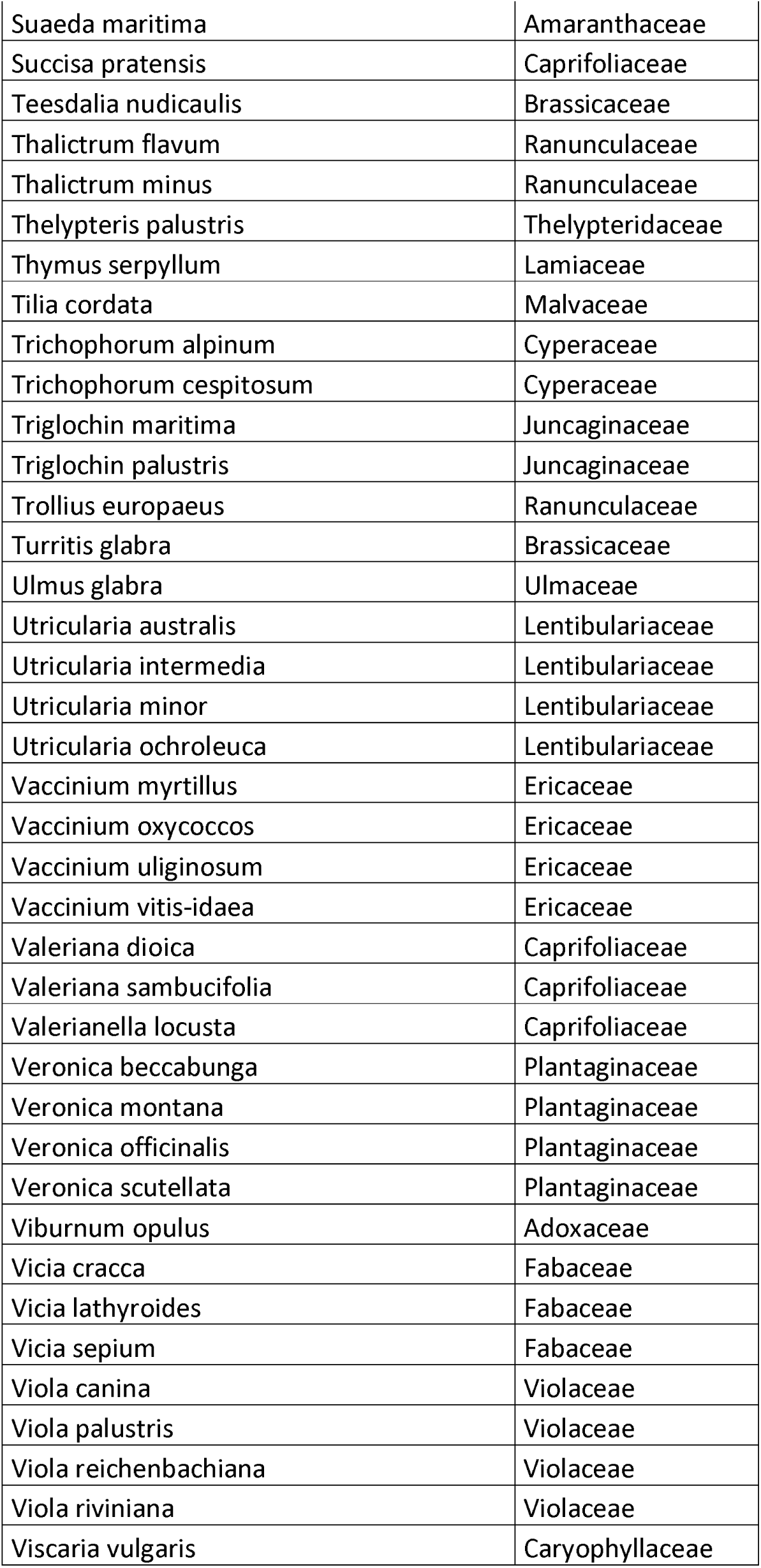

